# Mistargeting of secretory cargo in retromer-deficient cells

**DOI:** 10.1101/2020.07.02.185660

**Authors:** Sarah D. Neuman, Erica L. Terry, Jane E. Selegue, Amy T. Cavanagh, Arash Bashirullah

## Abstract

Intracellular trafficking is a basic and essential cellular function required for delivery of proteins to the appropriate subcellular destination; this process is especially demanding in professional secretory cells, which synthesize and secrete massive quantities of cargo proteins via regulated exocytosis. The *Drosophila* larval salivary glands are professional secretory cells that synthesize and secrete mucin proteins at the onset of metamorphosis. Using the larval salivary glands as a model system, we have identified a role for the highly conserved retromer complex in trafficking of secretory granule membrane proteins. We demonstrate that retromer-dependent trafficking via endosomal tubules is induced at the onset of secretory granule biogenesis, and that recycling via endosomal tubules is required for delivery of essential secretory granule membrane proteins to nascent granules. Without retromer function, nascent granules do not contain the proper membrane proteins; as a result, cargo from these defective granules is mistargeted to Rab7-positive endosomes, where it progressively accumulates to generate dramatically enlarged endosomes. Retromer complex dysfunction is strongly associated with Alzheimer’s disease, characterized by accumulation of amyloid β (Aβ). We show that amyloid precursor protein (APP) undergoes regulated exocytosis and accumulates within enlarged endosomes in retromer-deficient cells. These results highlight recycling of secretory granule membrane proteins as a critical step during secretory granule maturation and provide new insights into our understanding of retromer complex function in secretory cells. Our data also suggests that misrouting of secretory cargo, including APP, may contribute to the progressive nature of neurodegenerative disease.

**SUMMARY STATEMENT:** Retromer complex dysfunction is implicated in neurodegeneration. Here the authors show a new role for the retromer complex in recycling of secretory membrane and cargo proteins during regulated exocytosis.

## INTRODUCTION

A critical component of the regulated exocytosis machinery is the secretory granule; these specialized organelles contain cargo proteins surrounded by a phospholipid bilayer embedded with secretory granule membrane fusion proteins. Regulated exocytosis begins with the biogenesis of secretory granules at the *trans*-Golgi Network (TGN). Nascent secretory granules initially bud from the TGN as immature granules; these immature granules then undergo a maturation process that renders them competent for exocytosis (Kögel and Gerdes, 2010; Bonnemaison et al., 2013). The specific processes that occur during maturation vary by cell type; however, maturation frequently involves homotypic fusion of immature granules to generate larger granules of a defined size (Bonnemaison et al., 2013; Wendler et al., 2001; Urbé et al., 1998; Tooze et al., 1991) and refinement of secretory granule cargo and membrane content (Tooze and Tooze, 1986; Bonnemaison et al., 2013; Kögel and Gerdes, 2010). Once maturation is completed, secretory granules are retained in the cytoplasm until an appropriate stimulus triggers their release. The release of secretory granule cargo is driven by membrane fusion proteins, such as SNAREs and Synaptotagmins, that are present on both the secretory granule membrane and the plasma membrane (Burgoyne and Morgan, 2003; Hong, 2005). These proteins catalyze fusion of the two disparate membranes to allow secretory granule cargo release.

The retromer complex is a highly conserved regulator of retrograde trafficking. This pathway is used to retrieve specific transmembrane proteins from endosomes and recycle them back to the TGN, thereby preventing their degradation in lysosomes (van Weering et al., 2010; Seaman et al., 1998). Retromer is composed of two functional subcomplexes: the tubulation complex and the cargo-selective complex (CSC) (van Weering et al., 2010; Small and Petsko, 2015). The tubulation complex is composed of a heterodimer of two Bin/Ampiphysin/Rvs (BAR) domaincontaining sorting nexins (Vps5 and Vps17 in yeast; SNX1/2 and SNX5/6 in mammalian cells); these proteins are capable of inducing membrane deformations to form endosomal tubules (van Weering et al., 2010; Seaman et al., 1998; van Weering et al., 2012). The CSC, composed of a heterotrimer of VPS26, VPS29, and VPS35, recognizes specific transmembrane proteins and directs them into endosomal tubules (Bonifacino and Hurley, 2008). The CSC then recruits accessory proteins to drive scission of small retromer-studded vesicles containing the transmembrane cargo; these vesicles are transported back to the TGN (Cullen and Korswagen, 2012). The CSC can also work with a different sorting nexin, SNX3, to mediate retrograde trafficking of transmembrane proteins from endosomes in a tubule-independent manner (Harterink et al., 2011; Zhang et al., 2011; Burd and Cullen, 2014). CSC binding to endosomes requires interaction with both Rab5, an early endosome protein, and Rab7, a late endosome protein (Rojas et al., 2008). In this manner, the retromer complex retrieves specific transmembrane cargoes from endosomal compartments and recycles them back to the TGN via endosomal tubules. Work by us and others has suggested that secretory granule membrane fusion proteins may traffic through endosomal compartments (Neuman and Bashirullah, 2018; Burgess et al., 2012), raising the intriguing possibility that these proteins may undergo retromer-dependent retrieval and recycling.

The *Drosophila* larval salivary glands synthesize and secrete mucin-like “glue” proteins via regulated exocytosis at the onset of metamorphosis (Beckendorf and Kafatos, 1976; Korge, 1977). Biogenesis of these mucin-containing secretory granules begins at the mid-third instar larval transition, about 24 h before the onset of metamorphosis (Beckendorf and Kafatos, 1976; Biyasheva et al., 2001). Nascent granules mature via homotypic fusion, reaching a terminal size of 5-8 μm in diameter (Reynolds et al., 2019; Neuman and Bashirullah, 2018; Niemeyer and Schwarz, 2000). Mature granules then undergo steroid hormone-dependent exocytosis beginning about 4 h before the onset of metamorphosis (Kang et al., 2017; Biyasheva et al., 2001). After exocytosis, mucins are expelled out of the lumen of the salivary gland and onto the surface of the animal, allowing the prepupa to adhere to a surface during metamorphosis. Notably, the larval salivary glands complete just one single, developmentally regulated cycle of secretory granule biogenesis, maturation, and exocytosis over a 24 h period. Thus, by using fluorescently tagged mucins coupled with markers for secretory granule membrane proteins and other organelles, we can observe the temporal sequence of intracellular trafficking events that contribute to secretory granule maturation and exocytosis in living cells.

Here, we use the *Drosophila* larval salivary glands to uncover a role for the retromer complex in regulated exocytosis. Our data demonstrates that there is a developmentally regulated increase in endosomal tubulogenesis that coincides with the onset of secretory granule biogenesis. Importantly, nascent secretory granules in *Vps26* mutant cells lack the appropriate membrane proteins, and our data suggests that these secretory granule membrane proteins undergo retromer-dependent trafficking via endosomal tubules. In the absence of appropriate membrane proteins, cargo proteins are mistargeted to the endolysosomal system and aberrantly accumulate within dramatically enlarged Rab7-positive endosomes. Our data also indicates that transmembrane cargo proteins, like Amyloid Precursor Protein (APP), which undergoes regulated exocytosis in salivary gland cells, are similarly missorted to the endolysosomal system. Overall, these results highlight a new role for the retromer complex in maturation of secretory granules during regulated exocytosis and provide new insights into the etiology of known intracellular trafficking phenotypes in retromer-deficient cells.

## RESULTS

### Trafficking via endosomal tubules is developmentally regulated at the onset of cargo biogenesis

In the *Drosophila* larval salivary glands, biogenesis of mucins begins in the middle of the third larval instar, about 20 h prior to their release via regulated exocytosis (Fig. 1A). To begin to test if the retromer complex plays a role in this process, we used qPCR to measure mRNA expression levels of retromer complex components in dissected salivary glands isolated both before and after the onset of cargo biogenesis. The *Drosophila* genome contains a single ortholog of the cargo-selective complex (CSC) components *Vps26, Vps29*, and *Vps35*, as well as three retromer-associated sorting nexins (*Snx1, Snx3*, and *Snx6*) (Zhang et al., 2011; Wang et al., 2014). We observed a significant, ~2-fold increase in the expression levels of *Snx1* and *Snx6* and smaller but still significant increases in the expression levels of *Snx3, Vps26, Vps29*, and *Vps35* in glands after the onset of cargo biogenesis (Fig. 1B). Because *Snx1* and *Snx6* are BAR domaincontaining proteins that are both necessary and sufficient for endosomal tubulogenesis (van Weering et al., 2012), this result may indicate that there is increased trafficking via endosomal tubules in salivary glands after the onset of cargo biogenesis. To test this, we used the endosome and lysosome marker LAMP-GFP (Burgess et al., 2012; Wang et al., 2014) for live-cell time-lapse imaging to assess tubule formation. LAMP-GFP primarily localized in puncta in pre-biogenesis glands; these puncta appeared stable and little tubule activity was observed (Fig. 1C, Movie 1). However, in post-biogenesis glands, we observed a significant increase in both the appearance and movement of endolysosomal tubules (Fig. 1C, Movie 2), indicating that there is a developmentally regulated increase in endolysosomal tubular trafficking at the onset of cargo biogenesis.

**Figure 1.**
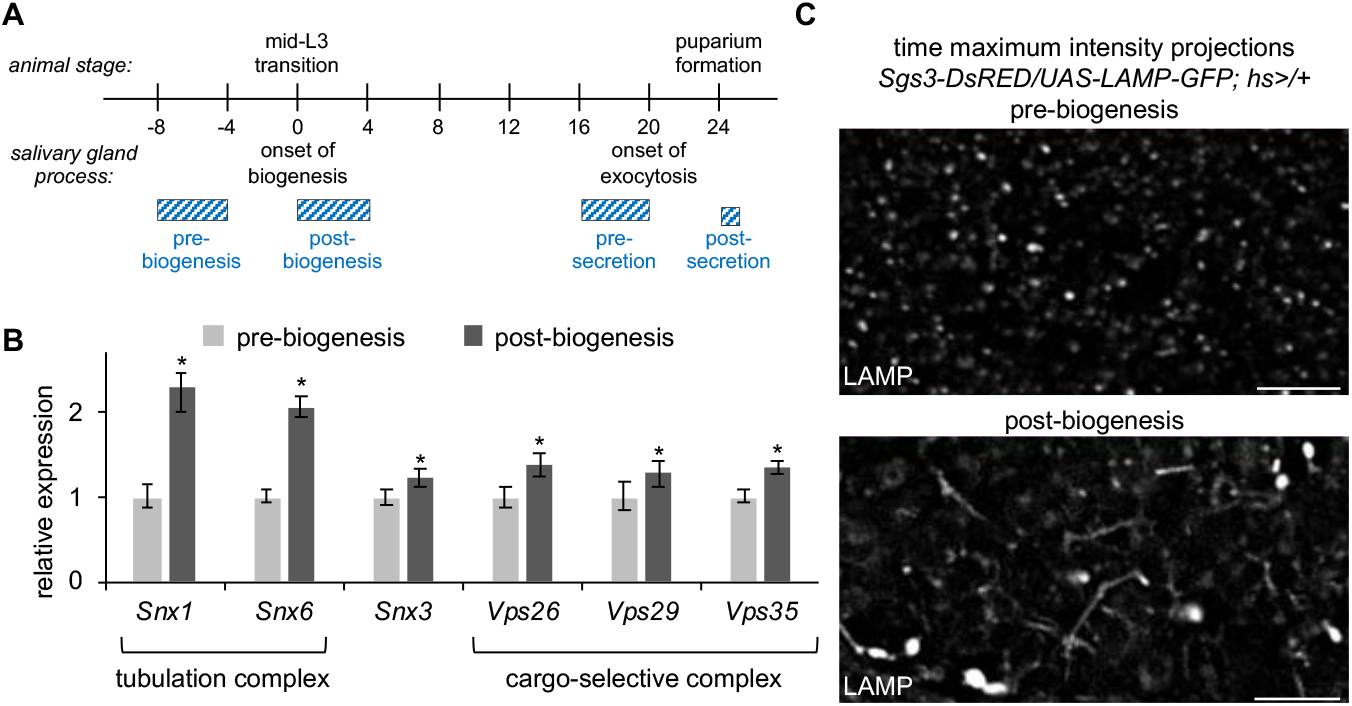
Endosomal tubule activity increases at the onset of secretory granule biogenesis. (A) Schematic depicting the timing of events during mucin regulated exocytosis in the larval salivary glands. Biogenesis of mucin secretory granules begins at the mid-third instar (mid-L3) larval transition. Exocytosis begins about 20 h later and is completed by puparium formation. Onset of biogenesis to completion of exocytosis spans 24 h. Developmental stages of salivary glands analyzed throughout this paper are depicted by the blue bars. (B) qPCR analysis of retromer complex mRNA expression levels pre- and post-biogenesis of secretory granules shows significant upregulation of all retromer complex genes in post-biogenesis salivary glands. *y*-axis shows relative expression; *x*-axis shows the genes analyzed. Samples were run in biological triplicate and normalized to the reference gene *rp49*. Error bars and statistics were calculated by REST analysis; asterisks indicate *p*<0.05. (C) Live-cell time-lapse imaging of LAMP-GFP shows an increase in tubulogenesis in salivary glands post-biogenesis. Data shown as time maximum intensity projections reflecting 250 timepoints (~3 min total time) acquired from a single optical slice. Scale bars 5 μm.

### Endosomal tubules traffic secretory granule membrane proteins to nascent granules

Defects in retromer complex function are strongly associated with neurodegenerative diseases, including Alzheimer’s and Parkinson’s disease (Wang and Bellen, 2015). The genetic lesions associated with these diseases primarily map to CSC components (Small et al., 2005; Wen et al., 2011; Muhammad et al., 2008; Zimprich et al., 2011; Vilariño-Güell et al., 2011). We therefore utilized loss of CSC function as an experimental context to explore the role of the retromer complex in regulated exocytosis. We found that endogenous Vps35-TagRFP protein (Koles et al., 2016) partially co-localized with endogenous Rab7-EYFP (Dunst et al., 2015) in control glands (Fig. S1A), but Vps35-TagRFP was absent in the salivary glands of loss-of-function *Vps26* mutant animals (*Vps26^B^*; Wang et al., 2014) (Fig. S1B), indicating that the CSC is non-functional in this mutant background. To determine if secretory granule biogenesis was affected by loss of retromer-dependent trafficking, we examined nascent secretory granules. Secretory granules are composed of two primary components: cargo proteins and membrane proteins. In the larval salivary glands, we can observe secretory cargo proteins via fluorescent tagging of mucins (*Sgs3-DsRED*). We saw that mucins were produced in *Vps26* mutant cells, and the nascent granules appeared similar to those of controls at the onset of biogenesis (Fig. 2A, B). Similarly, we can observe secretory granule membrane proteins via fluorescent tagging of Synaptotagmin-1 (Syt1-GFP) (Neuman and Bashirullah, 2018). All nascent granules had Syt1-GFP on their membrane in control cells (Fig. 2A). In contrast, we found that a significant number of nascent secretory granules in *Vps26* mutant cells did not have Syt1-GFP on their membrane (Fig. 2B), suggesting that there may be a defect in trafficking of secretory granule membrane proteins in retromer mutant cells.

**Figure 2.**
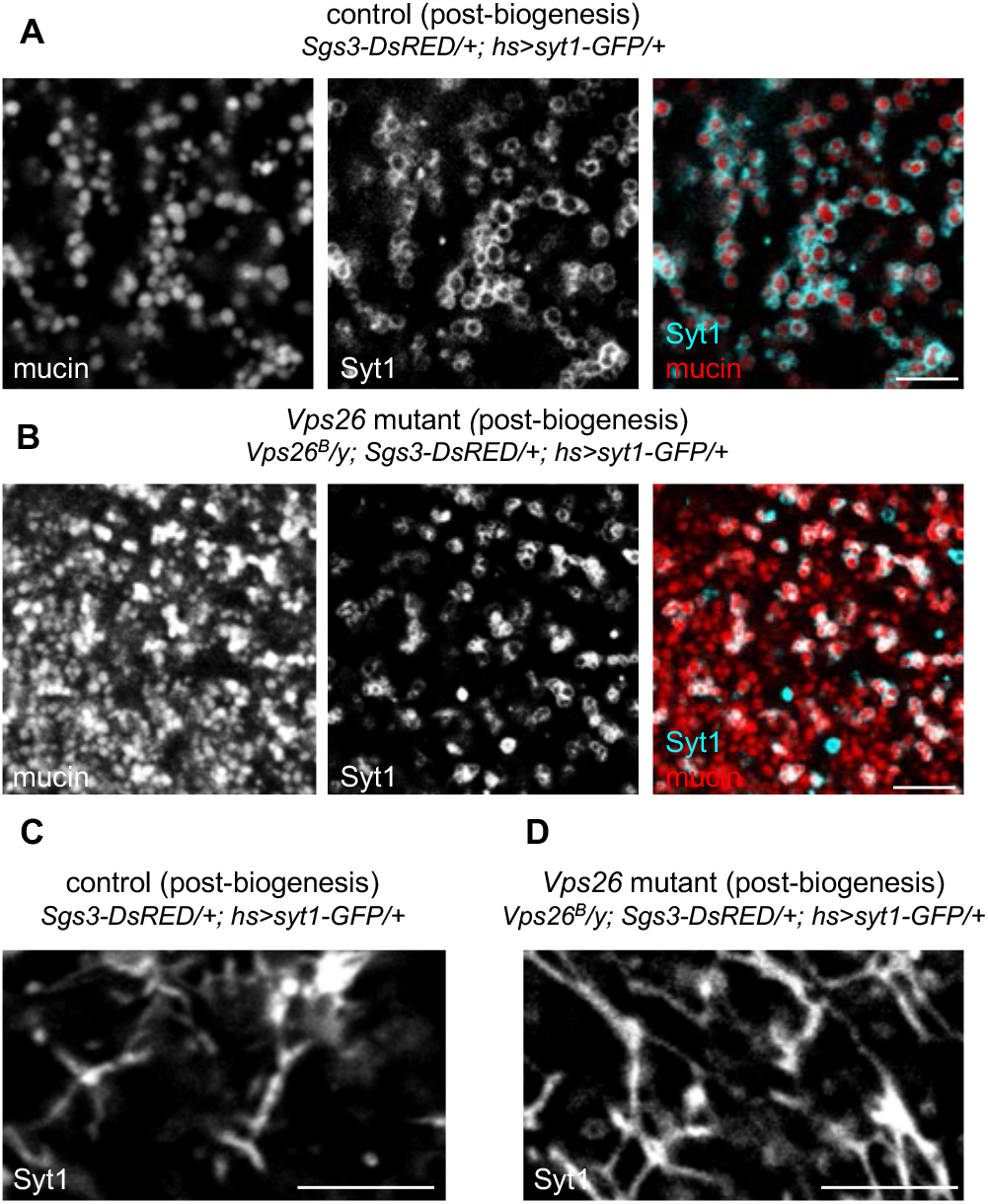
Secretory granule membrane protein trafficking is disrupted in retromer mutant cells. (A) Live-cell imaging of mucins (Sgs3-DsRED; red) and the secretory granule membrane protein Synaptotagmin-1-GFP (Syt1; cyan) in control salivary glands 0-4 h after the onset of secretory granule biogenesis shows that all nascent granules are surrounded by Syt1. (B) Live-cell imaging of mucins and Syt1 in *Vps26* mutant cells shows that a significant number of nascent granules lack Syt1 at biogenesis. (C) Live-cell imaging of Syt1 in control cells shows that Syt1 localizes in a tubular network. Note that the salivary glands are composed of polarized epithelial cells, and tubules are localized near the basal membrane; few secretory granules are visible in this plane. (D) Live-cell imaging of Syt1 in *Vps26* mutant cells shows that Syt1 localizes in a more extensive tubular network in retromer-deficient cells. Scale bars 5 μm.

Work by us and others has suggested that secretory granule membrane proteins may traffic through the endosomal system (Neuman and Bashirullah, 2018; Burgess et al., 2012). We therefore wanted to test if Syt1-GFP trafficked through endosomal tubules. Strikingly, we saw that Syt1-GFP localized in a tubular network that closely resembled that of LAMP-GFP in post-biogenesis control cells (Fig. 2C). We also observed an expansion of this Syt1-GFP tubular network in *Vps26* mutant cells (Fig. 2D). Because SNX1 protein expression has been reported to increase in VPS26 mutant cells (Seaman, 2004), we would expect to see more endosomal tubules in the absence of CSC function. Taken together, these results suggest that secretory granule membrane proteins undergo retromer-dependent trafficking through the endolysosomal network, and this trafficking is required for delivery of membrane proteins to nascent secretory granules.

### Nascent granules lacking appropriate membrane proteins become enlarged and are not secreted

Secretory granule membrane proteins, including Synaptotagmins and SNAREs, are essential for fusion with the plasma membrane during regulated exocytosis. Given that many secretory granules lacked Syt1-GFP at biogenesis in *Vps26* mutant cells, we tested if there were any regulated exocytosis defects in these cells. Live cell imaging of mucins at puparium formation, the developmental stage when all mucins should be secreted, revealed that a great deal of cargo was retained in the salivary gland cells of *Vps26^B^* mutant animals (Fig. 3A). *Vps26* mutant *flp/FRT* somatic clones in the salivary glands also displayed retention of mucins at prepupal stages (Fig. 3B), indicating that a considerable amount of mucins are never secreted and that these defects are cell-autonomous.

**Figure 3.**
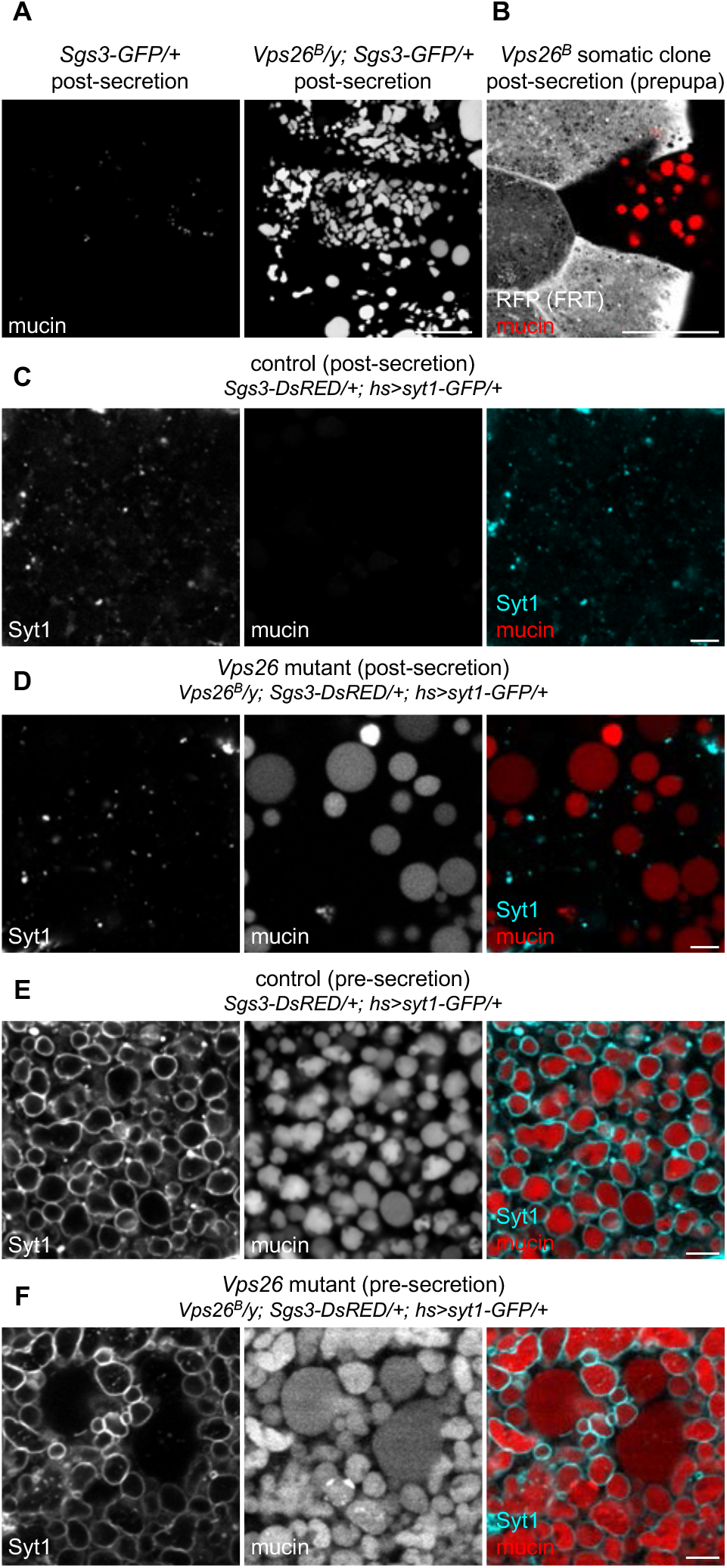
Regulated exocytosis of secretory cargo is disrupted in retromer mutant cells. (A) Live-cell imaging of mucins (Sgs3-GFP) in control and *Vps26* mutant salivary glands post-secretion. All mucin cargo proteins are secreted in control cells, while *Vps26* mutant cells still contain large amounts of cargo. (B) *Vps26^B^* somatic clone shows that mucin secretion defects are cell-autonomous. The mutant clone is marked by loss of RFP, shown in gray, and mucins are shown in red. Note that the mutant clone is the only cell still containing mucins in prepupal salivary glands. Full genotype: *Vps26^B^, FRT19A/hs-flp, UbiRFPnls, FRT19A; Sgs3-GFP*/+. (C) Live-cell imaging of mucins (Sgs3-DsRED; red) and Syt1-GFP (cyan) in control salivary gland cells post-secretion. Mucins have been secreted, and Syt1 localizes in small puncta within the cytoplasm. (D) Live-cell imaging of mucins and Syt1 in *Vps26* mutant cells shows that a significant number of mucin-containing secretory granules are not secreted; these granules are also enlarged and lack Syt1 on their membrane. (E) Live-cell imaging of mucins and Syt1 in control cells pre-secretion. All mucin-containing secretory granules have Syt1 on their membrane, and granules are of a relatively uniform size. (F) Live-cell imaging of mucins and Syt1 in *Vps26* mutant cells pre-secretion. Many normal-sized mucin-containing secretory granules have Syt1 on their membrane; however, enlarged mucin granules lack Syt1. Scale bars in A, B are 50 μm; scale bars in C-F are 5 μm.

Importantly, imaging of *Vps26* mutant salivary glands at puparium formation (post-secretion) demonstrated that retained mucin granules lacked Syt1-GFP on their membranes (Fig. 3C, D). These granules also appeared to be significantly enlarged compared to the maximum size of mucin granules in control cells pre-secretion (*c.f*. Fig. 3D vs. 3E). We also analyzed the morphology and membrane protein content of granules immediately prior to secretion, when maturation is complete and granules have reached their final size. At this stage in control cells, all cargo-containing secretory granules had Syt1-GFP on their membrane and were of a relatively uniform size (Fig. 3E); however, in *Vps26* mutant cells, we observed a population of enlarged, mucin-containing vesicles that did not have Syt1-GFP on their membrane, although the remaining granules of normal size did have Syt1-GFP (Fig. 3F). Therefore, our results thus far demonstrate that two distinct populations of mucin granules develop in retromer-deficient cells. First is a population of granules that have appropriate membrane proteins at biogenesis; these granules achieve normal size and are secreted. Second is a population of granules that lack the appropriate membrane proteins at biogenesis; here, mucins accumulate in enlarged compartments that are not secreted.

### Secretory cargo accumulates in enlarged Rab7-positive endosomes

Because homotypic fusion during maturation requires secretory granule membrane proteins, the nascent granules lacking Syt1-GFP in *Vps26* mutant cells should not be able to undergo this process. Therefore, the dramatically enlarged granules observed just prior to and after secretion must reach this size by other mechanisms. Because the retromer complex is known to regulate retrograde trafficking from Rab5/Rab7-positive endosomes (Rojas et al., 2008), we first analyzed the localization and morphology of these endosomes using endogenously-regulated, EYFP-tagged Rab5 and Rab7 (Dunst et al., 2015). There were no significant differences in the localization or morphology of Rab5-positive early endosomes in *Vps26* mutant cells (Fig. S2A); however, Rab7-positive late endosomes were dramatically enlarged and often misshapen in salivary gland cells from *Vps26* mutant animals and in *Vps26* mutant *flp/FRT* somatic clones (Fig. S2B, C). Strikingly, these enlarged, Rab7-positive endosomes contained mucins in both pre- and post-secretion *Vps26* mutant salivary gland cells (Fig. 4B, D), while mucins were not seen inside Rab7-positive endosomes in control cells (Fig. 4A, C). To confirm these results, we also assessed the localization of mucins with a GFP-tagged FYVE domain (FYVE-GFP), which binds to the lipid phosphatidylinositol-3-phosphate (PI3P) present in endosomal membranes (Wucherpfennig et al., 2003). Consistent with our Rab7 results, we observed enlarged, mucin-containing vesicles wrapped in FYVE-GFP in *Vps26* mutant glands both pre- and post-secretion (Fig. S3). Taken together, these results suggest that cargo proteins accumulate in the endosomal system in retromer mutant cells.

**Figure 4.**
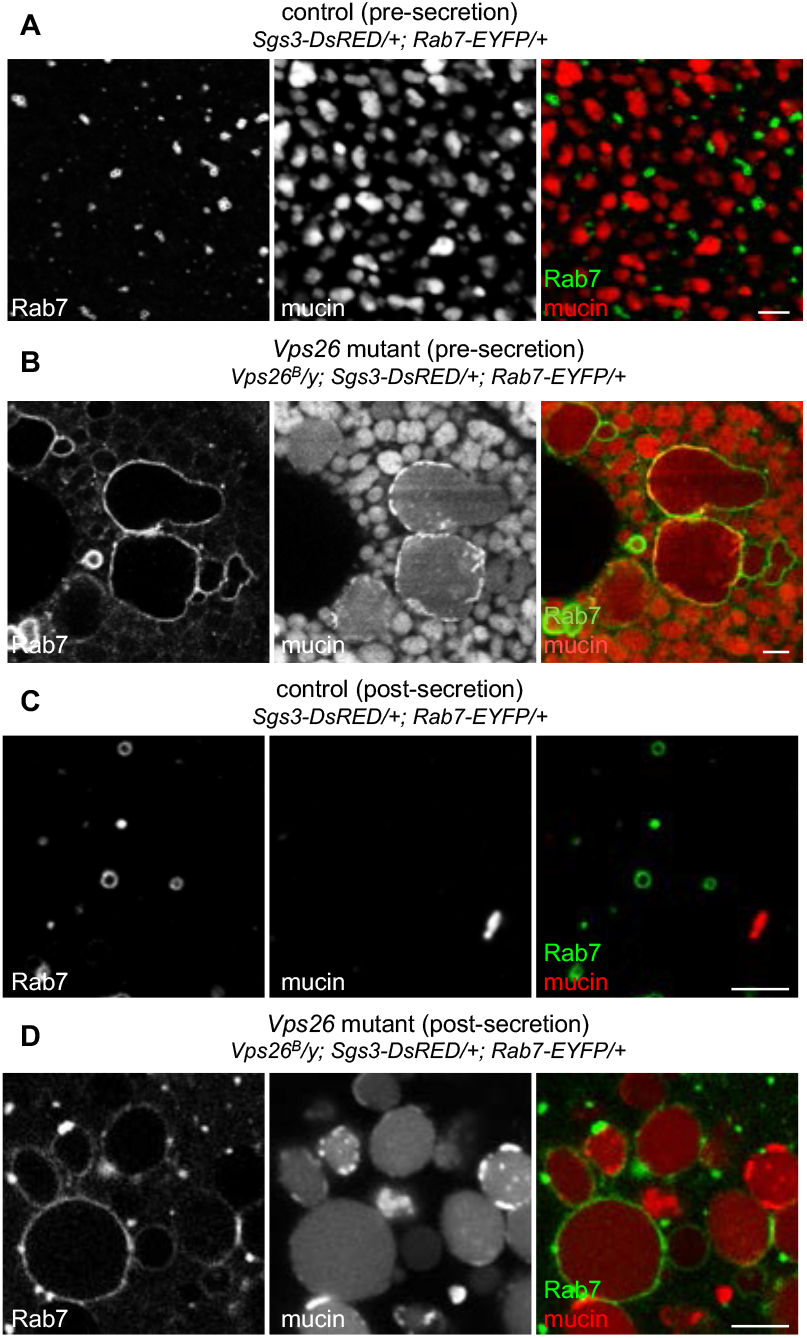
Enlarged Rab7-positive endosomes contain secretory cargo proteins in retromer mutant cells. (A) Live-cell imaging of Rab7-EYFP (green) and mucins (Sgs3-DsRED; red) in control salivary glands pre-secretion. Rab7 localizes in small puncta that are completely distinct from mucins. (B) Live-cell imaging of Rab7 and mucins in *Vps26* mutant salivary glands pre-secretion. Note that Rab7-positive endosomes are dramatically enlarged and contain mucin cargo proteins. (C) Live-cell imaging of Rab7 and mucins in control salivary glands post-secretion. Most mucins have been secreted and Rab7 localizes in small vesicles. (D) Live-cell imaging of Rab7 and mucins in *Vps26* mutant salivary glands post-secretion. Unsecreted mucin granules are dramatically enlarged and surrounded by Rab7. Scale bars 5 μm.

### Mistargeting of secretory cargo proteins to endosomes begins immediately following secretory granule biogenesis

Our next goal was to determine when endosomal accumulation of mucins begins in *Vps26* mutant cells. The larval salivary glands enable us to answer this question because we can observe the entire process of regulated exocytosis in real time. Because we observed Syt1-GFP trafficking defects at secretory granule biogenesis, we tested if mucin trafficking was also affected at this stage. In control cells, Rab7 localized in small puncta that were completely distinct from nascent granules during secretory granule biogenesis (Fig. 5A). In contrast, *Vps26* mutant cells exhibited enlarged, Rab7-positive vesicles which contained mucins at this same developmental stage (Fig. 5B). To confirm these results, we also analyzed the localization of FYVE-GFP with mucin cargo proteins. Like Rab7, FYVE-GFP localized in puncta that were distinct from nascent secretory granules in control cells (Fig. 5C), while *Vps26* mutant cells contained mucins surrounded by FYVE-GFP (Fig. 5D). Interestingly, these mucin-containing, Rab7- and FYVE-GFP-positive endosomes were only present ~4 h after the onset of biogenesis, while mucins lacking Syt1-GFP were visible at the onset of biogenesis, suggesting that mucin cargo proteins lacking Syt1-GFP are taken up by endosomes after nascent granules have budded from the TGN.

**Figure 5.**
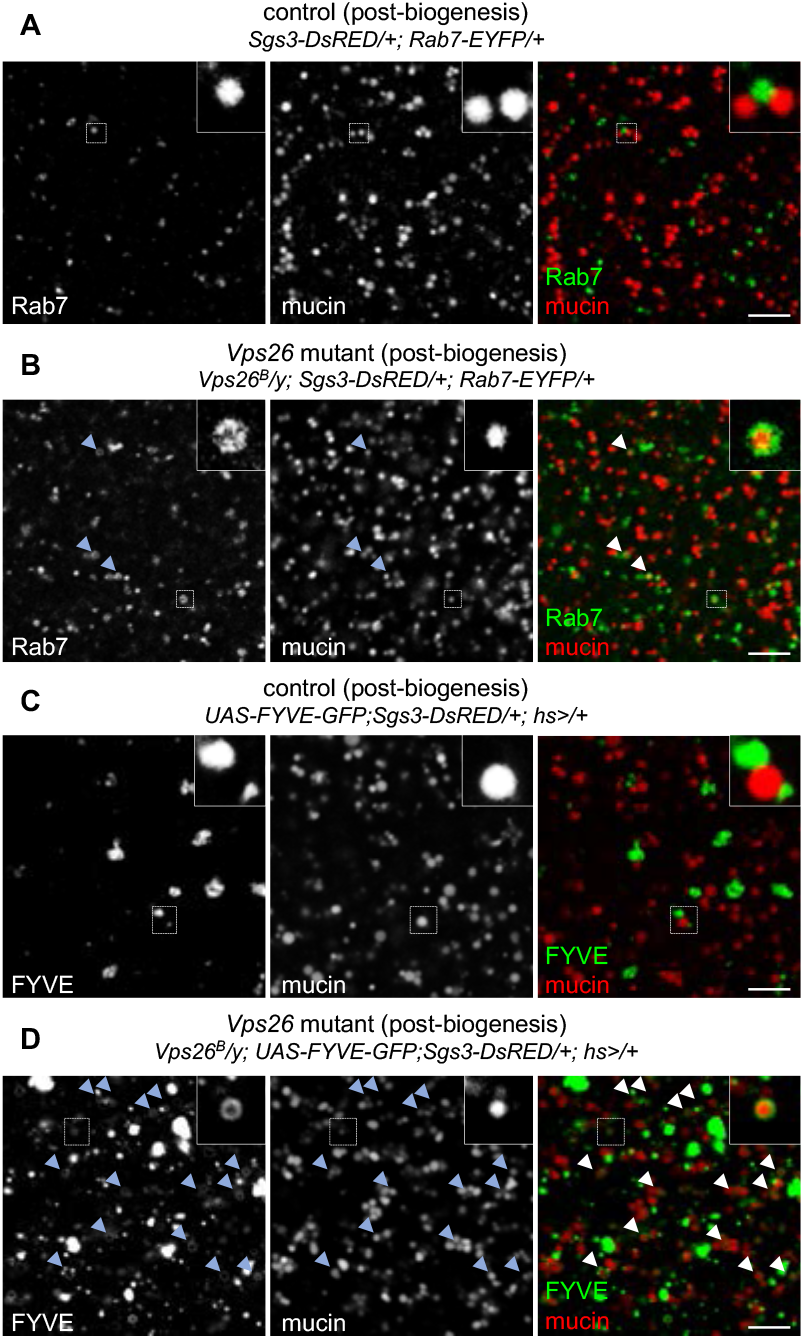
Secretory cargo begins to accumulate within endosomes at biogenesis. (A) Live-cell imaging of Rab7-EYFP (green) and mucins (Sgs3-DsRED; red) in control cells immediately following the onset of secretory granule biogenesis. Rab7 localizes in small puncta that are completely distinct from mucins. Boxed area is magnified at the upper right. (B) Live-cell imaging of Rab7 and mucins in *Vps26* mutant cells immediately following the onset of secretory granule biogenesis. Rab7 endosomes appear slightly enlarged compared to controls (A) and some contain mucin cargo, indicated by arrowheads. Boxed region is magnified at upper right. (C) Live-cell imaging of the PI3P marker FYVE-GFP (green) and mucins (Sgs3-DsRED; red) in control salivary glands post-biogenesis. FYVE-GFP and mucins are completely distinct; boxed region magnified at upper right. (D) Live-cell imaging of FYVE-GFP and mucins in *Vps26* mutant salivary glands post-biogenesis. A significant number of mucins are surrounded by FYVE-GFP, indicated by arrowheads. Boxed region is magnified at upper right. Scale bars 5 μm.

The retromer complex recycles the cation-independent mannose-6 phosphate receptor (CI-M6PR), a carrier protein for lysosomal hydrolases (Seaman, 2004; Arighi et al., 2004). Without recycling of CI-M6PR, lysosomal hydrolases are not efficiently delivered, resulting in lysosomal dysfunction (McMillan et al., 2017). To test whether lysosomal dysfunction results in accumulation of mucin cargo proteins within endosomes, we looked for retention of mucins in loss-of-function *Lerp* (the fly ortholog of CI-M6PR) mutant cells (Hasanagic et al., 2015). Importantly, *Lerp* mutant salivary glands did not contain any mucin cargo proteins at the onset of metamorphosis (Fig. S4B), indicating that there are no secretion defects in this mutant background. To confirm that *Lerp* mutant salivary glands did have lysosomal dysfunction, we used LysoTracker staining to label acidified organelles. Control glands at puparium formation contained many small LysoTracker-positive puncta, consistent with the expected size of acidified late endosomes and lysosomes (Fig. S4A). In contrast, *Lerp* mutant salivary glands contained enlarged LysoTracker-positive organelles that appeared dense in a DIC image (Fig. S4B). These phenotypes confirm that there is lysosomal dysfunction in *Lerp* mutant glands. Overall, this data suggests that lysosomal dysfunction is not the cause of aberrant mucin cargo protein accumulation in retromer mutant cells; instead, our data suggests that mucins are mistargeted to the endosomal system upon loss of retromer complex function.

### Membrane-bound cargo proteins, like APP, are also mistargeted to the endolysosomal system in retromer mutant cells

Defects in retromer complex function are strongly associated with neurodegenerative diseases, including Alzheimer’s disease (Small et al., 2005; Wen et al., 2011; Wang and Bellen, 2015; Muhammad et al., 2008). One of the hallmarks of Alzheimer’s disease is the accumulation of amyloid-β (Aβ) plaques (Stelzmann et al., 1995); Aβ is a cleavage product of endosome-associated processing of Aβ Precursor Protein (APP) (Reiss et al., 2018), a membrane-bound, secreted protein that is known to undergo retromer-dependent retrograde trafficking via recycling of the Sortilin-LA (SorLA) sorting receptor (Fjorback et al., 2012; Lane et al., 2012; Wang and Bellen, 2015). In Alzheimer’s disease, it is thought that retromer loss-of-function results in failure to recycle SorLA, resulting in retention of APP in endosomes, where APP can be processed by β- and γ-secretase to generate aberrantly large quantities of Aβ (Small and Gandy, 2006). However, our data suggests that there may be additional mechanisms that lead to endosomal accumulation of APP. First, surprisingly, we found that ectopically expressed APP is secreted via regulated exocytosis in salivary gland cells. Regulated exocytosis of mucin granules occurs at the apical membrane of the glands (Rousso et al., 2015; Tran et al., 2015; Biyasheva et al., 2001), and we observed that APP was absent from the apical membrane but present on mucin granule membranes in control cells prior to the onset of secretion (Fig. 6A, B). However, after the onset of mucin secretion, APP was present on the apical membrane (Fig. 6B), indicating that it had been deposited there during regulated exocytosis. We then examined the localization of APP in *Vps26* mutant cells. Pre-secretion, we found that APP was still present on secretory granule membranes; however, it was also present in large clumps that co-localized with LysoTracker staining (Fig. 6C), indicating that these are likely enlarged, acidified late endosomes or lysosomes. These results demonstrate that APP is normally secreted to the plasma membrane via regulated exocytosis in control salivary gland cells. However, in retromer mutant cells, APP accumulates in the endolysosomal system prior to its appearance on the plasma membrane. This data suggests that APP is directly trafficked to the endolysosomal system in retromer-deficient cells. Taken together, these results suggest that both soluble and membranebound secretory cargo proteins are mistargeted to endosomes in retromer mutant cells.

**Figure 6.**
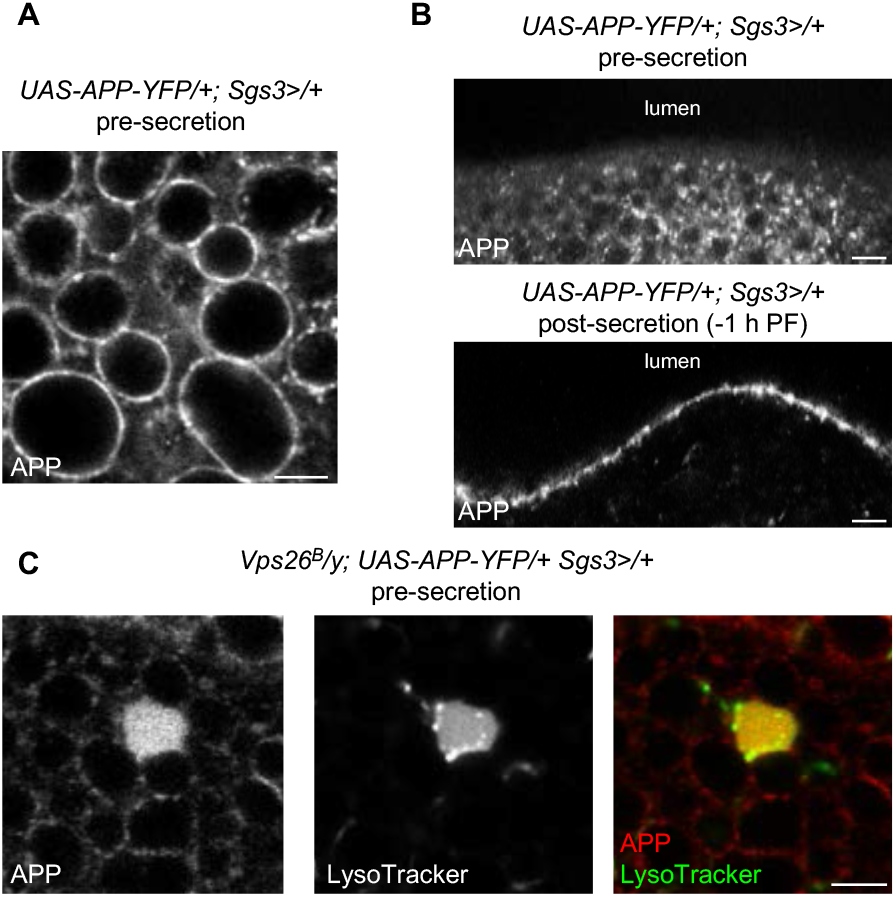
APP undergoes regulated exocytosis and is missorted to endosomes in retromer mutant cells. (A) Live-cell imaging of APP-YFP in control salivary glands pre-secretion. Note that APP is enriched on mucin secretory granule membranes. (B) Live-cell imaging of APP localization in control glands pre- and post-secretion. The apical membrane of the salivary glands faces the lumen (top of images). APP is absent from the apical membrane pre-secretion but is enriched on the apical membrane post-secretion, indicating that APP undergoes regulated exocytosis in salivary gland cells. (C) Live-cell imaging of APP (red) and LysoTracker (green) to label acidified organelles in *Vps26* mutant cells. APP accumulates in enlarged, LysoTracker-positive compartments in *Vps26* mutant cells pre-secretion. Scale bars 5 μm.

## DISCUSSION

Professional secretory cells synthesize massive quantities of cargo proteins that are destined for regulated exocytosis. Therefore, it is imperative that these cells exercise careful control over the delivery of proteins to the appropriate subcellular locations. Here, we have identified a role for the retromer complex in recycling of secretory granule membrane proteins to nascent granules; delivery of these membrane proteins promotes secretory granule maturation and competence for secretion. Failure to deliver secretory granule membrane proteins to nascent granules results in the mistargeting and progressive accumulation of secretory cargo proteins within endosomes. These results reveal new insights into our understanding of retromer loss-of-function phenotypes and their role in disease etiology.

One of the critical steps of secretory granule maturation in many different cell types is the homotypic fusion of immature granules to generate larger, mature granules of a defined size (Bonnemaison et al., 2013; Kögel and Gerdes, 2010). These membrane fusion events are mediated by secretory granule membrane proteins, like Synaptotagmins and SNAREs; therefore, the presence of these proteins on nascent granules is obligatory for maturation to proceed. Synaptotagmins and SNAREs are also required later for secretory granule fusion with the plasma membrane during regulated exocytosis. Currently, it is thought that both secretory granule membrane proteins and secretory granule cargo proteins are produced via *de novo* synthesis; these proteins are translated in the rough endoplasmic reticulum (ER), trafficked through the Golgi cisternae, and packaged into nascent granules via budding from the TGN. Our results partially support this model, as we observe that some nascent secretory granules do possess the appropriate membrane proteins in retromer-deficient cells; these granules appear to mature normally and are eventually secreted. However, we also observe that many nascent granules lack the proper membrane proteins in retromer mutant cells, suggesting that there is an alternative, retromer-dependent pathway for delivery of secretory granule membrane proteins. In this alternative pathway, secretory granule membrane proteins are recycled through endosomal compartments to nascent granules. Our data suggests that this retromer-dependent recycling pathway is critical during regulated exocytosis, as a significant fraction of nascent secretory granules rely on it to acquire the proper membrane proteins during maturation.

Although our data suggests that secretory granule membrane proteins are new cargo proteins for the retromer complex, the mechanism by which retromer recognizes these proteins remains unclear. Typically, the Vps26-Vps29-Vps35 heterotrimer that composes the cargo-selective complex, is, as the name implies, responsible for identification of transmembrane cargo proteins that undergo retrograde trafficking/recycling. Identification and capture of cargo has typically been reported to require interaction between VPS35 and specific motifs on the cytoplasmic tail of transmembrane proteins (Seaman, 2004; Arighi et al., 2004; Nothwehr et al., 1999; Seaman, 2007; Fjorback et al., 2012; Mukadam and Seaman, 2015). However, no specific “consensus” sorting motif for CSC binding has been identified. Our data suggests that Synaptotagmin-1, and possibly other secretory granule membrane proteins, are new retromer cargo proteins. Future studies will be required to identify the mechanism by which the retromer complex identifies secretory granule membrane proteins, including analysis of potential sorting motifs.

Retromer complex dysfunction has frequently been reported to result in enlarged endosomes (Wang et al., 2014; Arighi et al., 2004; Maruzs et al., 2015). This defect has largely been attributed to failure to remove retromer cargo proteins from endosomes or deemed a passive consequence of impaired lysosomal function resulting from reduced delivery of lysosomal hydrolases. However, our results suggest an alternative explanation. We observe that secretory cargo proteins progressively accumulate within endosomal compartments, resulting in endosomes of enormous size. Additionally, we do not observe accumulation of secretory cargo proteins when lysosomal function is disrupted by loss of the lysosomal hydrolase carrier. Therefore, we propose that mistargeting of secretory cargo is a novel etiological factor in the development of enlarged endosomes in retromer-deficient cells.

Endosomal dysfunction is also strongly associated with neurodegenerative disease. Indeed, early studies of Alzheimer’s disease found that enlarged endosomal compartments were almost completely diagnostic of the disease (Cataldo et al., 2000). The brains of Alzheimer’s patients were found to be deficient in the CSC components VPS26 and VPS35 (Small et al., 2005), and loss of CSC function resulted in prolonged retention of APP within endosomes, where APP was more likely to undergo amyloidogenic processing to generate abnormally large quantities of Aβ (Berman et al., 2015; Small and Gandy, 2006). The retention of APP within endosomes has largely been tied to defects in retromer-dependent retrograde recycling of VPS10 family members, like SorLA, which act as sorting receptors for APP, or to defects in retromer-dependent recycling of APP itself (Small and Petsko, 2015; Berman et al., 2015). However, our data suggests an alternative model. We observe that APP is secreted via regulated exocytosis, and it accumulates in enlarged, acidified compartments prior to the onset of secretion in retromer-deficient cells. Overall, our data suggests that missorting of secretory cargo proteins may contribute to endosomal dysfunction, shedding new light on the etiology and progressive nature of neurodegenerative disease.

## MATERIALS AND METHODS

### Drosophila stocks, husbandry, somatic clones, and developmental staging

The following fly stocks were obtained from the Bloomington *Drosophila* Stock Center: *w^1118^, Sgs3-GFP, Rab5-EYFP, Rab7-EYFP, Sgs3-GAL4, hs-GAL4, UAS-GFP-LAMP, UAS-GFP-myc-2xFYVE, UAS-APP-YFP, UAS-Synaptotagmin-1-GFP, Vps26^B^, Vps35-TagRFP, hs-flp, UbiRFPnls, FRT19A*, and *Lerp^F6^. Sgs3-DsRED* was provided by A. Andres (University of Nevada, Las Vegas, NV, USA). Note that animals from experiments using *hs-GAL4* were not heat-shocked; the *hs-GAL4* transgenic construct ‘leaks’ constitutively in the salivary glands due to a cryptic promoter in the transgene (Costantino et al., 2008). All experimental crosses were grown on standard cornmeal-molasses media in uncrowded vials or bottles and maintained at 25°C. For *Vps26^B^* somatic clones in the salivary glands, embryos were heat-shocked at 37°C for 30 min 0-4 h after egg lay. Staged animals collected pre-secretion were selected as wandering L3 (wL3) larvae at about −8 h PF (~4 h before the onset of secretion) and imaged under a fluorescent dissecting microscope to ensure that no glue protein was present in the lumen. Animals collected post-secretion were staged as white prepupae (0 h PF) for the onset of puparium formation. For analyses pre- and post-mucin biogenesis, animals were collected 0-4 h after hatching from the embryo (0-4 h after L1) and allowed to age at 25°C for the appropriate time. Animals pre-biogenesis were collected at 56-60 h after L1 (8 h prior to the induction of glue biosynthesis). For 0-4 h after biogenesis, early L3 (eL3) larvae were collected and analyzed under a fluorescent dissecting microscope to confirm the absence of *Sgs3-GFP* or *Sgs3-DsRED* expression. Animals lacking mucin expression were then allowed to age for 4 h at 25°C, then were again analyzed for *Sgs3-GFP* or *Sgs3-DsRED* expression. Animals expressing the transgene at this time were collected for analysis.

### Quantitative real-time PCR (qPCR)

qPCR was performed as previously described (Ihry et al., 2012). Total RNA was isolated from salivary glands at the appropriate developmental stage (see staging details in section above) using the RNeasy Plus Mini Kit (Qiagen). cDNA was synthesized from 400 ng of total RNA using the SuperScript III First Strand Synthesis System (Invitrogen). Samples were collected and analyzed in biological triplicate, and minus reverse transcriptase controls were conducted to confirm the absence of genomic DNA contamination. qPCR was performed using a Roche LightCycler 480 with LightCycler 480 SYBR Green I Master Mix (Roche). Amplification efficiencies were calculated for each primer pair and minus template controls were included for each experiment. Relative Expression Software Tool (REST) was used to calculate relative expression and *p*-values (Pfaffl et al., 2002). Expression was normalized to the reference gene *rp49*; these primers were previously published (Ihry et al., 2012). Retromer complex gene primers were designed using FlyPrimerBank (Hu et al., 2013); primer sequences are listed in Table S1.

### Confocal microscopy

All images were obtained from live, unfixed salivary glands; at least 10 glands were imaged per experiment. Imaging was carried out at room temperature. Salivary glands were dissected in PBS from animals of the appropriate genotype and developmental stage and mounted in either PBS or 1% low melt agarose (Apex Chemicals); glands were imaged for no longer than 10 min after mounting. Images were acquired using either an Olympus FV1000 laser scanning microscope (40x oil immersion objective, NA 1.30; 60x oil immersion objective, NA 1.42; 100x oil immersion objective, NA 1.40) with FV10-ASW software or an Olympus FV3000 laser scanning confocal microscope (100x oil immersion lens, NA 1.49) with FV31S-SW software. Brightness and contrast were adjusted post-acquisition using either FV10-ASW or FV31S-SW software. Images in Fig. S1 were collected from 3 optical sections at a 0.35 μm step size, comprising a total of 0.7 μm depth. These images were then deconvolved using three iterations of the Olympus CellSens Deconvolution for Laser Scanning Confocal Advanced Maximum Likelihood algorithm. The images displayed in the figure show a single optical slice. Images in Fig. 1 C, D and Movie 1, 2 were collected as live time-lapse movies using the Olympus FV3000 resonant scan head. Three optical sections at a 0.36 μm step size (total depth 0.72 μm) were collected at a frame rate of 1.37 frames/sec for Fig. 1C pre-biogenesis/Movie 1 and 1.40 frames/sec for Fig. 1C post-biogenesis/Movie 2. 250 frames were collected for each sample. Images were deconvolved as described above. The images in Fig. 1 C represent a time maximum intensity projection for a single optical slice, while Movies 1 and 2 display a maximum intensity projection of 3 optical slices played over time. For LysoTracker staining, LysoTracker Deep Red (Invitrogen) was diluted 1:1000 in PBS, and dissected salivary glands were incubated in this solution for 1 min on a shaker platform. Stained glands were rinsed for 1 min in PBS, then mounted and imaged immediately.

## ACKNOWLEDGEMENTS

Stocks obtained from the Bloomington *Drosophila* Stock Center (NIH P40OD018537) were used in this study. This work was supported in part by the National Institutes of Health (GM123204 to A.B.).

## AUTHOR CONTRIBUTIONS

Conceptualization: S.D.N., A.B; Methodology: S.D.N., A.B.; Validation: S.D.N., A.B.; Investigation: S.D.N., E.L.T., J.E.S., A.T.C.; Data curation: S.D.N.; Writing-original draft: S.D.N.; Writing-review and editing: S.D.N., A.B.; Visualization: S.D.N., A.B.; Supervision: A.B.; Project administration: A.B.; Funding acquisition: A.B.

---

**Figure S1.**
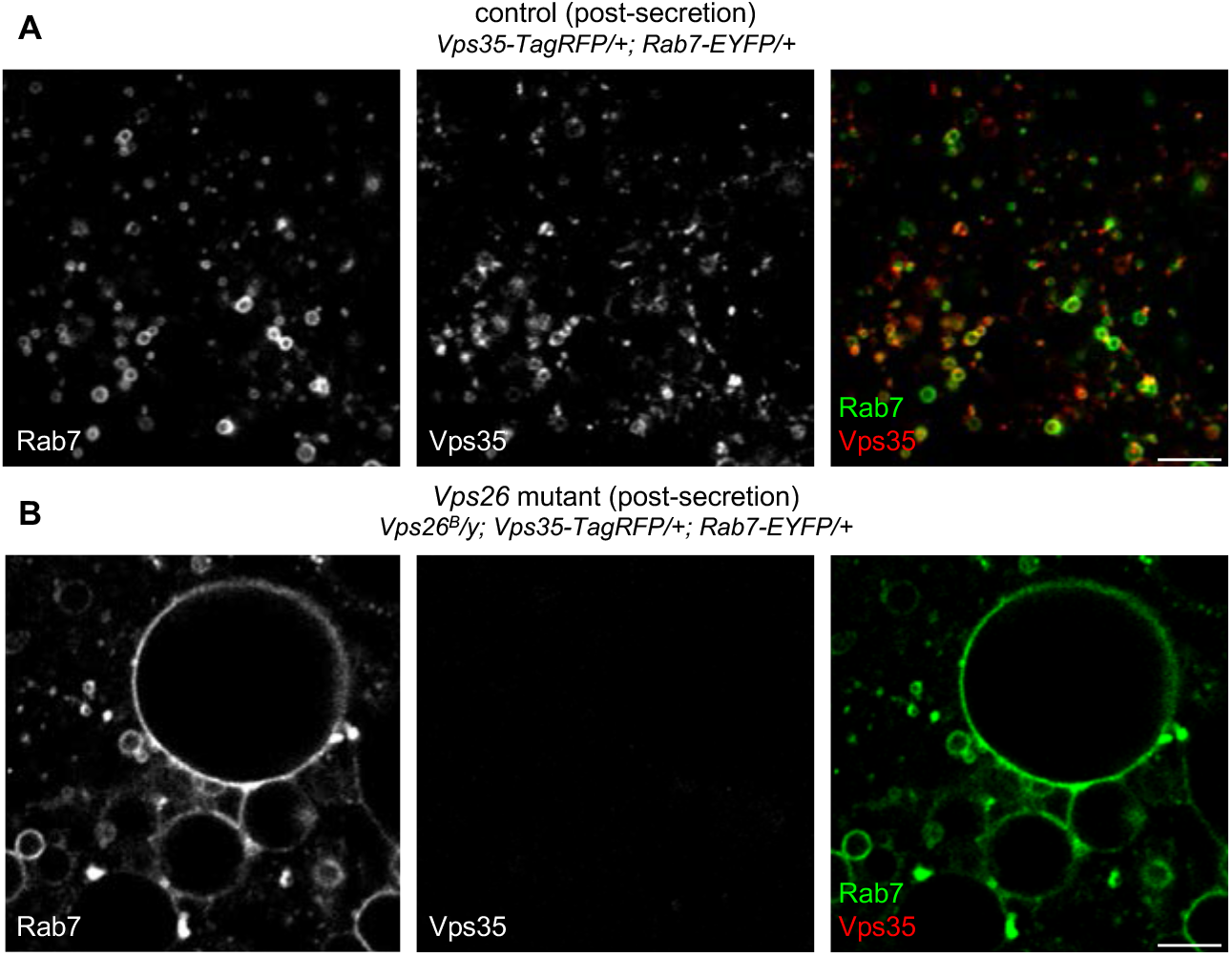
Vps35 protein is absent in *Vps26* mutant salivary gland cells. (A) Live-cell imaging of Rab7-EYFP (green) and Vps35-TagRFP (red) in control salivary gland cells post-secretion. As expected, Vps35 partially co-localizes with Rab7. (B) Live-cell imaging of Rab7 and Vps35 in *Vps26* mutant salivary gland cells post-secretion. Vps35 protein expression is undetectable, and Rab7-positive endosomes are significantly enlarged in *Vps26* mutant cells (see Fig. 4 and Fig. S2). Images were taken using identical laser power and acquisition settings. Scale bars 5 μm.

**Figure S2.**
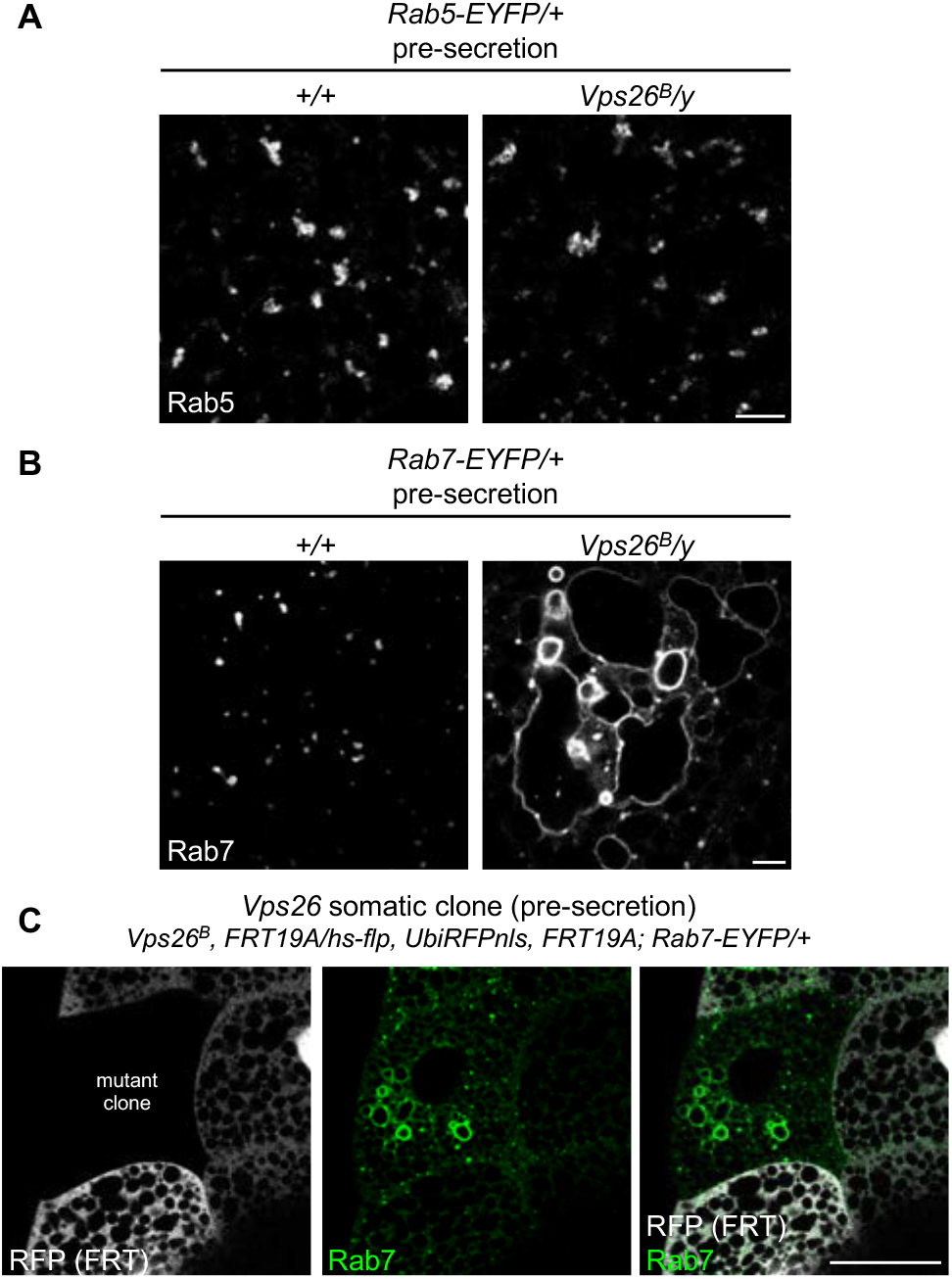
Analysis of endosomal morphology in *Vps26* mutant cells. (A) Live-cell imaging of Rab5-EYFP in control and *Vps26* mutant cells shows that the morphology of Rab5-positive endosomes appears to be unaffected in *Vps26* mutant cells. (B) Live-cell imaging of Rab7-EFYP in control and *Vps26* mutant cells shows that Rab7-positive endosomes are dramatically enlarged in *Vps26* mutant cells. (C) Somatic *Vps26* mutant clones show that Rab7 endosomal morphology defects are cell-autonomous. The mutant clone is marked by loss of RFP (shown in gray); Rab7 shown in green. Only the mutant clone contains enlarged Rab7-positive endosomes. Note that most endosomes are localized near the basal membrane in salivary gland cells, while the enlarged endosomes are found in the medial region of the cell. Scale bars in A, B are 5 μm; scale bar in C is 50 μm.

**Figure S3.**
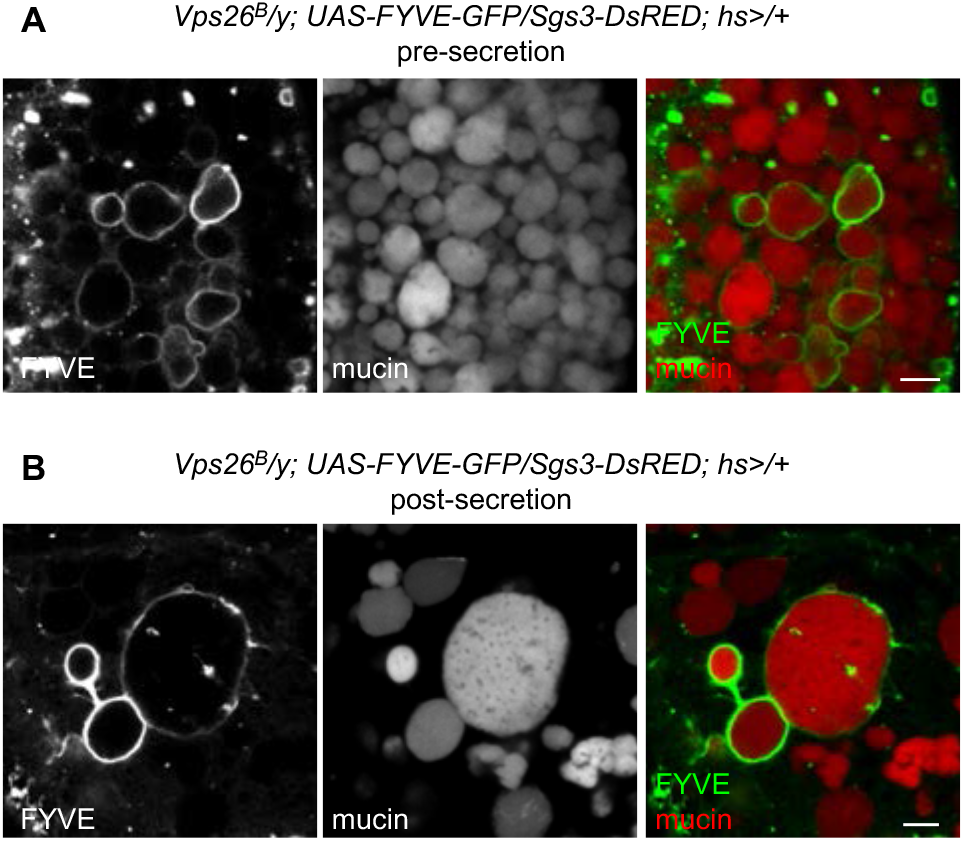
Enlarged mucin granules are surrounded by FYVE-GFP in *Vps26* mutant cells. Live-cell imaging of the PI3P-binding marker FYVE-GFP (green) and mucins (Sgs3-DsRED; red) in *Vps26* mutant cells pre- (A) and post-secretion (B). At both developmental stages, enlarged mucin granules are wrapped in FYVE-GFP. Scale bars 5 μm.

**Figure S4.**
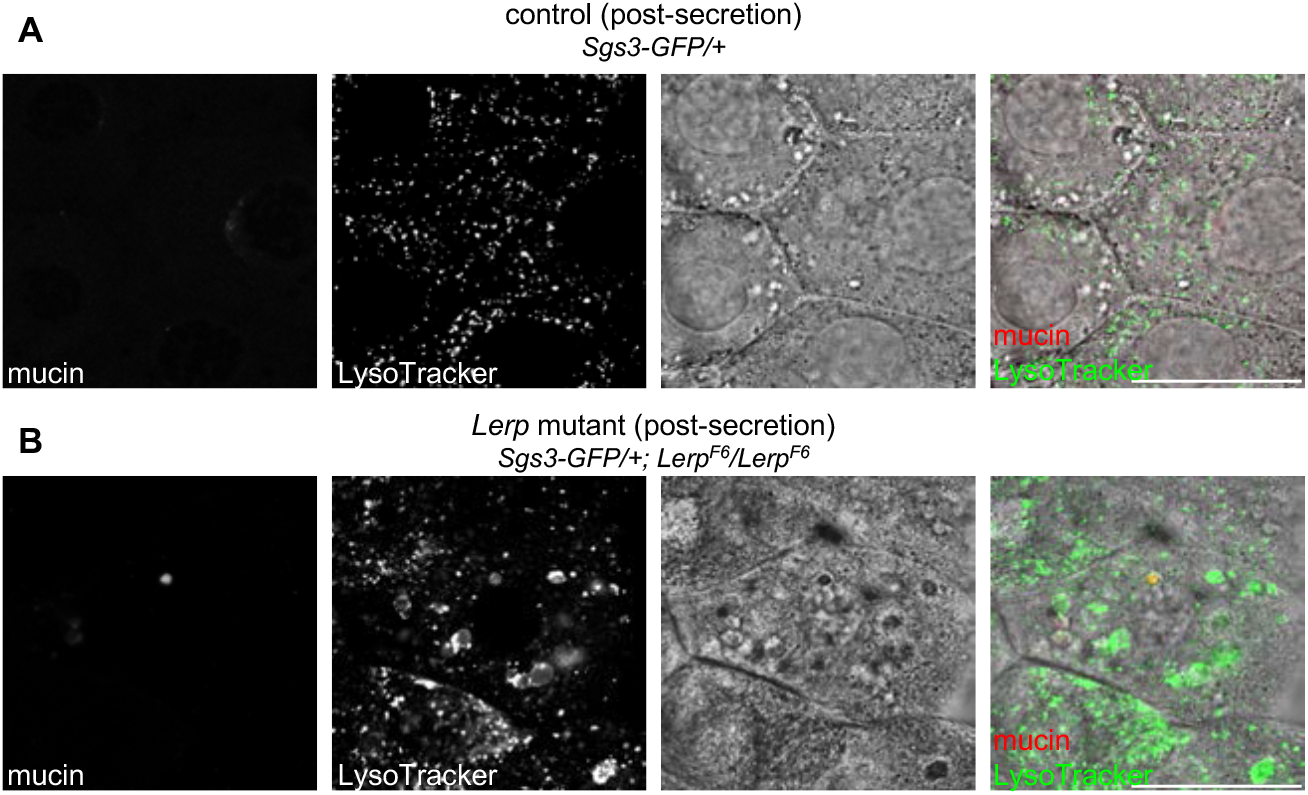
Lysosomal dysfunction does not cause mistargeting of secretory cargo proteins. (A) Live-cell imaging of mucins (Sgs3-GFP; red) and LysoTracker (green) in control cells post-secretion. All mucins have been secreted, and LysoTracker localizes in small puncta consistent with the size of acidified late endosomes and lysosomes. DIC image shows normal cellular morphology. (B) Live-cell imaging of mucins and LysoTracker in *Lerp* (fly CI-M6PR/lysosomal hydrolase carrier) mutant cells post-secretion. Mucins have been secreted, but many enlarged, LysoTracker-positive compartments are present. These enlarged, acidified compartments appear dark on a DIC image, suggesting the presence of undegraded material in dysfunctional, enlarged lysosomes and endosomes. Scale bars 50 μm.

**Supplemental Movie 1. Time-lapse imaging of endosomal tubule activity in pre-biogenesis salivary glands.** Live-cell time-lapse imaging of LAMP-GFP in salivary glands prior to the onset of glue biogenesis shows little tubule activity. Movie shows a maximum intensity projection of three optical slices; movie plays at a frame rate of 30 frames/sec. Scale bar 5 μm. A time-intensity projection of this movie is displayed in Fig. 1C.

**Supplemental Movie 2. Time-lapse imaging of endosomal tubule activity in post-biogenesis salivary glands.** Live-cell time-lapse imaging of LAMP-GFP in salivary glands after the onset of glue biogenesis shows the presence of dynamic endosomal tubules. Movie shows a maximum intensity projection of three optical slices; movie plays at a frame rate of 30 frames/sec. Scale bar 5 μm. A time-intensity projection of this movie is displayed in Fig. 1C.

**Table S1:**
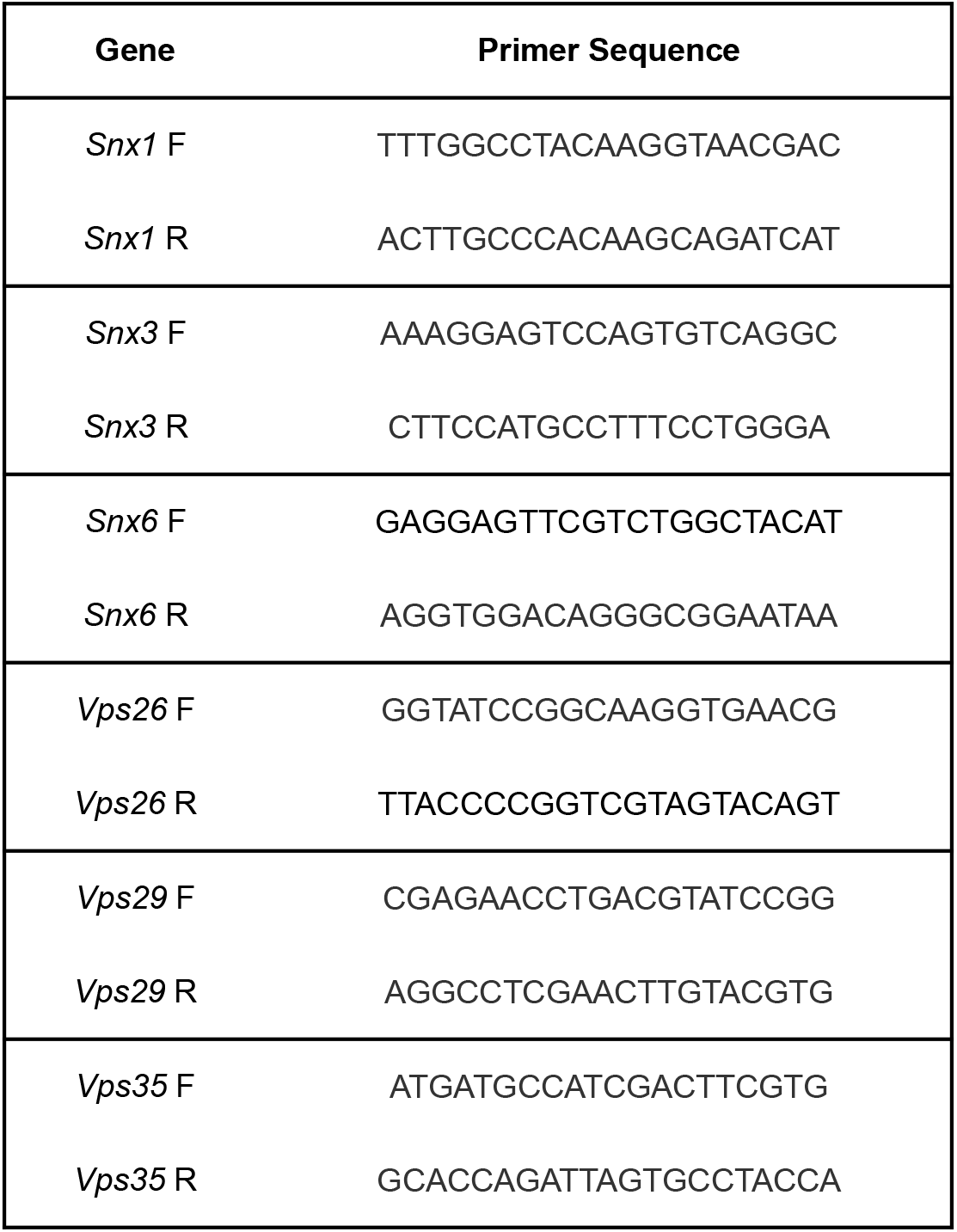
Primer sequences for qPCR

